## Extended Data Figures 1-7 for "CRL4^DCAF12^ regulation of MCMBP ensures optimal licensing of DNA replication"

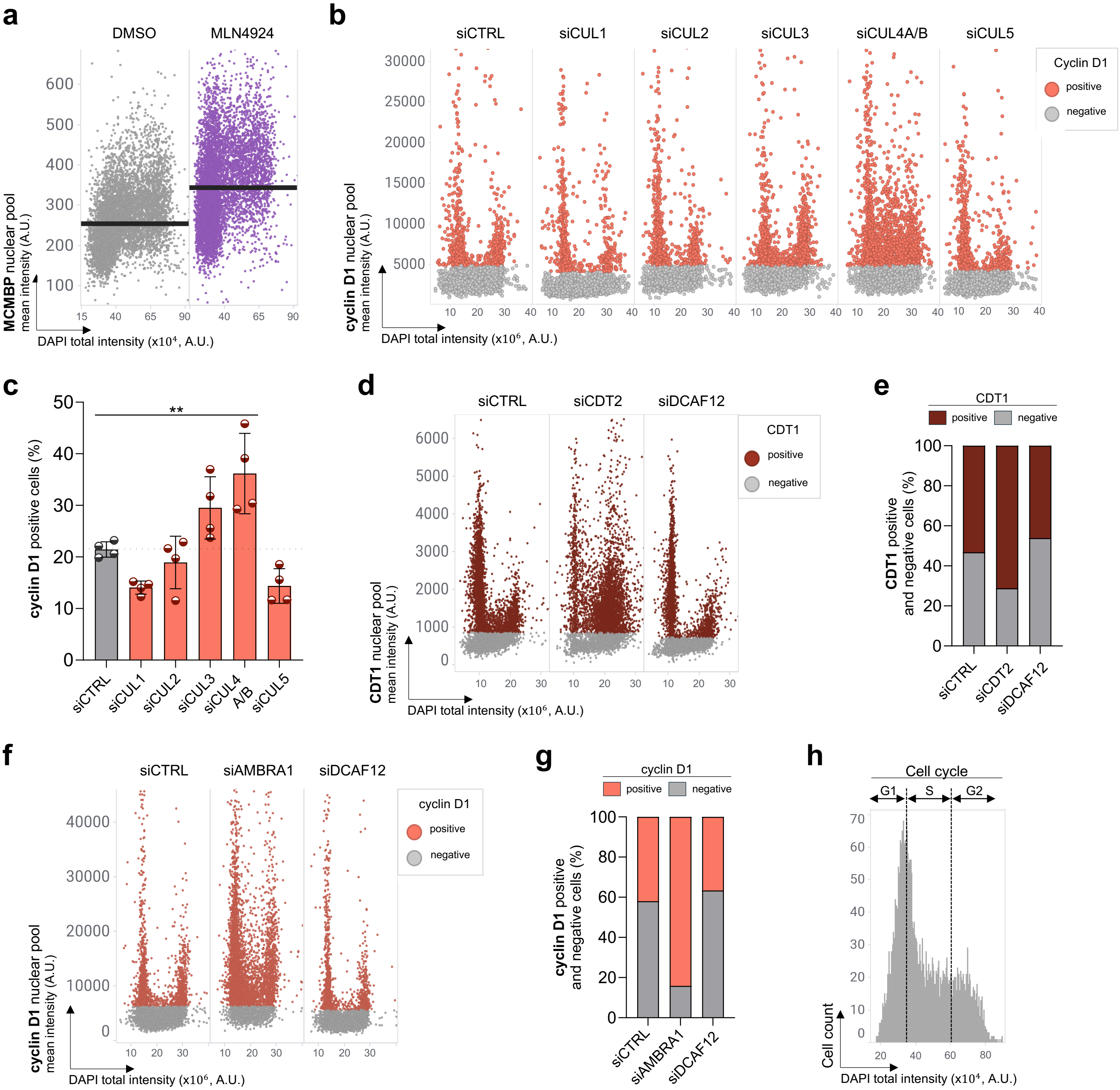


**Extended Data Fig. 1**

**Extended Data Fig. 1: MLN4924, a protein neddylation inhibitor, increases cellular levels of MCMBP.** **a,** QIBC plots of total nuclear MCMBP in U2OS cells after treatment with DMSO or MLN4924 (5 µM, 6 h). Lines denote medians; n ≈ 6,800 cells per condition. **b,** QIBC plots of MCM4-Halo cells immunostained for cyclin D1 after indicated siRNA treatments; n ≈ 5,000 cells per condition. **c,** Quantification of cyclin D1 positive cells (based on sub-stratification in **b**). Each bar indicates the percentage of cyclin D1 positive cells (data are mean ± s.d.; n = 4 technical replicates). **d,** QIBC plots of MCM4-Halo cells immunostained for CDT1 after indicated siRNA treatments; n ≈ 5,000 cells per condition. **e,** Quantification of QIBC plots in **d**; each bar indicates the percentage distribution of CDT1 positive and negative cells (based on sub-stratification in **d**). **f,** QIBC of MCM4-Halo cells immunostained for cyclin D1 after indicated siRNA treatments; n ≈ 5,000 cells per condition. **g,** Quantification of QIBC plots in **f**; each bar indicates the percentage distribution of cyclin D1 positive and negative cells (based on sub-stratification in **f**). **h,** DAPI profile-based sub-stratification of cell cycle phases in MCM4-Halo cells. P values were calculated by ordinary one-way ANOVA with Dunnett's test (**c**); $**P<0.01$. (A.U. Arbitrary Units).

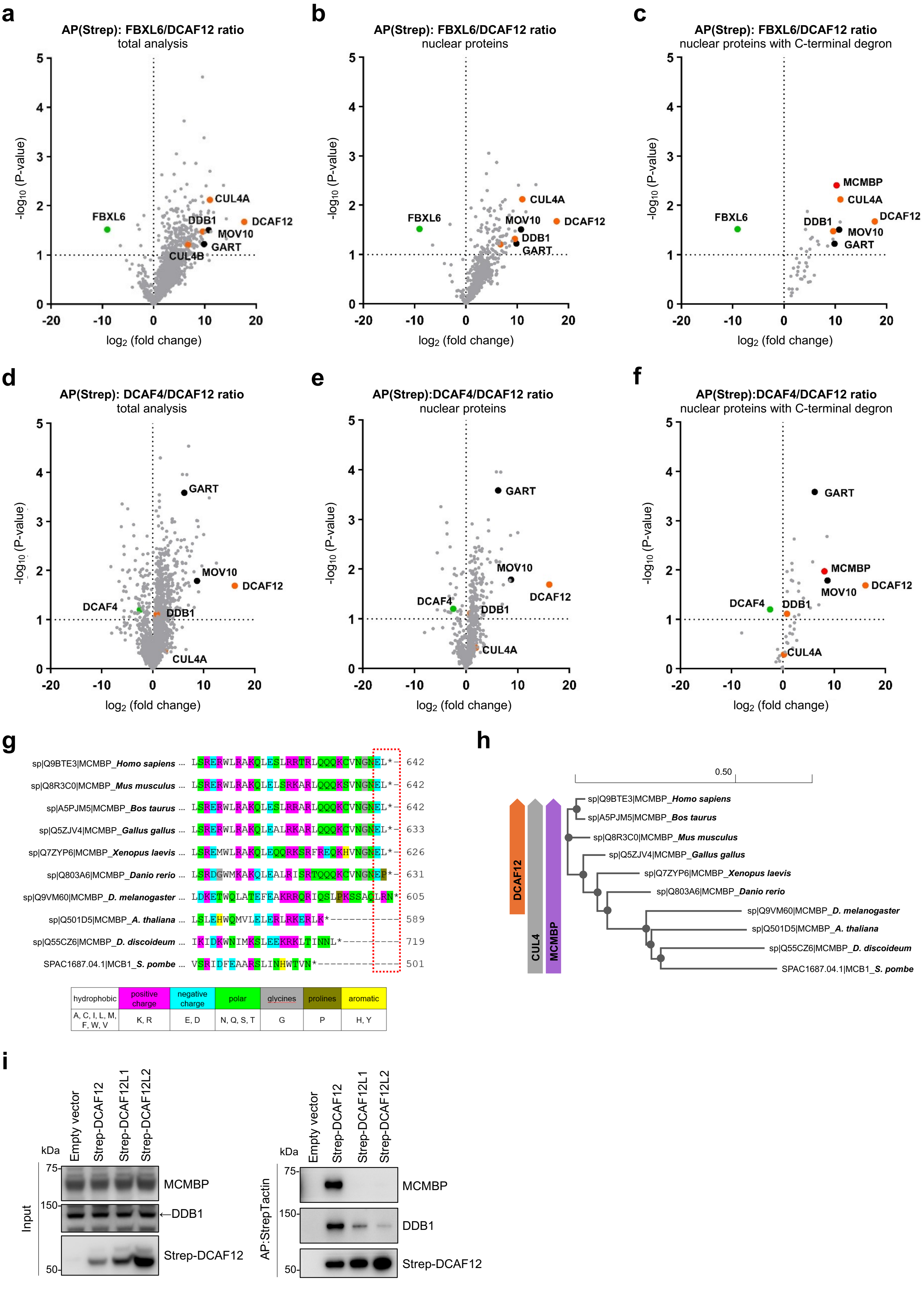


**Extended Data Fig. 2**

**Extended Data Fig. 2: MCMBP specifically interacts with DCAF12 but not with its mammalian paralogs.** **a,** $CRL4^{DCAF12}$-associated proteins identified by liquid chromatography-tandem mass spectrometry (LC-MS/MS). The composition of purified complexes from both steps was analyzed by LC-MS/MS. Non-related ubiquitin ligase $SCF^{FBXL6}$ was used as a control. Proteins that were significantly enriched with $CRL4^{DCAF12}$ compared to $SCF^{FBXL6}$ are in the upper right quadrant. **b,** Same as in a, but only nuclear proteins are shown. **c,** Same as in a, but only nuclear proteins with C-terminal acidic degron are shown. **d,** $CRL4^{DCAF12}$-associated proteins identified by liquid chromatography-tandem mass spectrometry (LC-MS/MS). The composition of purified complexes from both steps was analyzed by LC-MS/MS. Non-related ubiquitin ligase $CRL4^{DCAF4}$ was used as a control. Proteins that were significantly enriched with $CRL4^{DCAF12}$ compared to $CRL4^{DCAF4}$ are in the upper right quadrant. **e,** Same as in a, but only nuclear proteins are shown **f,** Same as in a, but only nuclear proteins with C-terminal acidic degron are shown. **g,** Multiple sequence alignment of the C-terminal sequence of MCMBP protein from indicated species. Sequences were aligned using EMBL-EBI Clustal Omega[45]. **h,** Phylogram of species used in multiple sequence alignment in **g** with indicated emergence of *DCAF12*, *CUL4*, and *MCMBP* genes. **i,** Affinity purification of N-terminal STREP-FLAG-tagged DCAF12 and its paralogs. HEK293T cells were co-transfected with StrepII-FLAG-tagged DCAF12, DCAF12L1 and DCAF12L2. MLN4924 was added for the final 6 hours before harvest. Following lysis, whole-cell lysates (WCL) were subjected to affinity purification (AP) with Strep-TactinXT resin and immunoblotted as indicated. The left panel shows 1% input samples, the right panel shows Strep-Tactin AP.

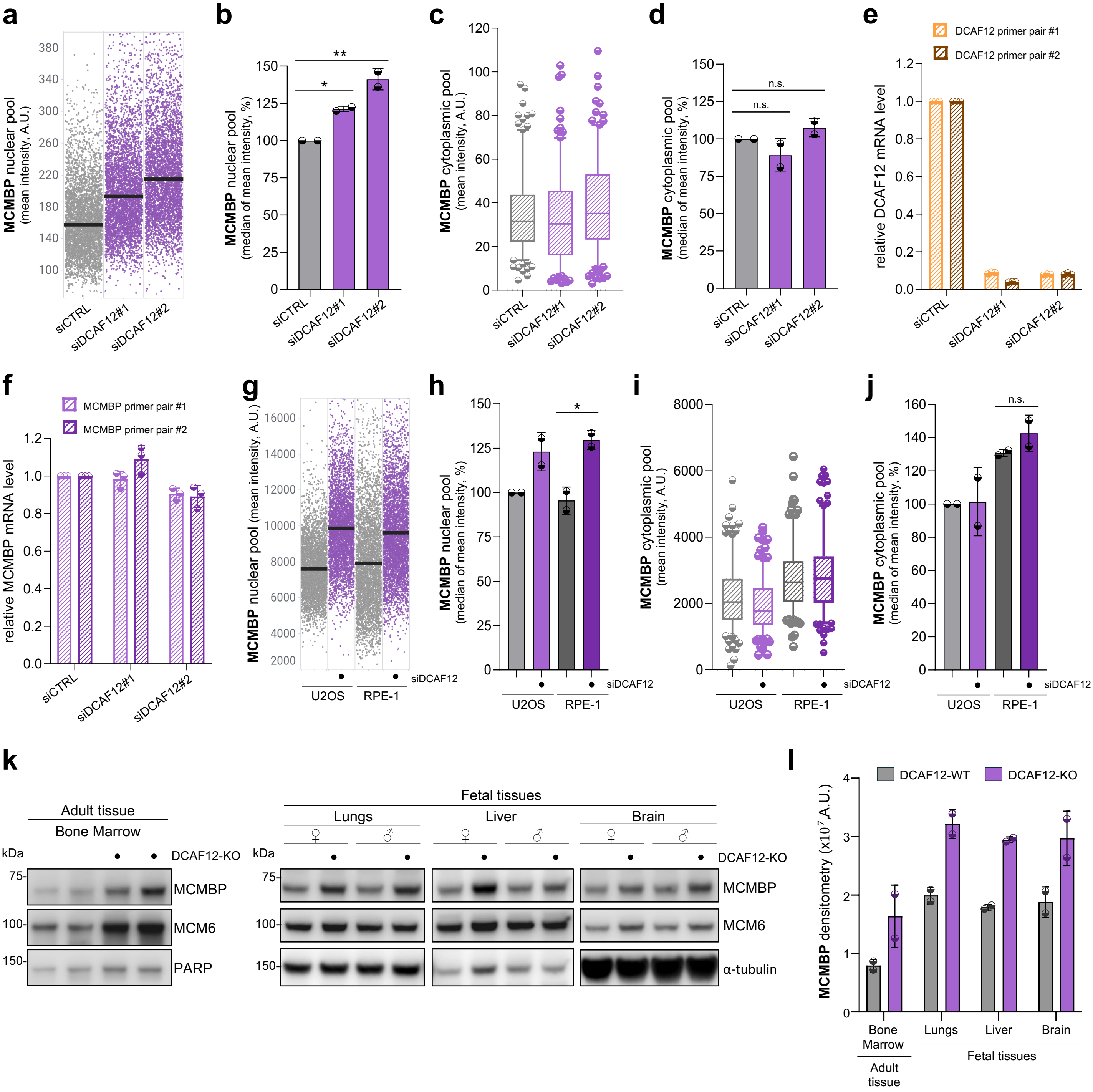


Extended Data Fig. 3

**Extended Data Fig. 3: Depletion of DCAF12 leads to stabilization of MCMBP protein levels in the nucleus. a,** QIBC plots of MCM4-Halo cells immunostained for MCMBP after treatment with control siRNA or siRNAs targeting different regions of the *DCAF12* gene (siDCAF12#1 or siDCAF12#2), as indicated. Lines denote medians; n ≈ 4,300 cells per condition. **b,** Quantification of QIBC plots in **a**, each bar indicates the median of mean intensity normalized with respect to siCTRL as 100 percent (data are mean ± s.d.; n = 2 biological replicates). **c,** MFI of cytoplasmic MCMBP after indicated siRNA treatments. **d,** Quantification of MFI of cytoplasmic MCMBP in **c**; each bar indicates the median of mean intensity normalized with respect to siCTRL as 100 percent (data are mean ± s.d.; n = 2 biological replicates). **e,** qPCR-based analysis of DCAF12 mRNA levels after indicated siRNA treatments; normalized with respect to siCTRL (data are mean ± s.d.; n = 3 technical replicates). **f,** qPCR-based analysis of MCMBP mRNA levels upon indicated siRNA treatments; normalized with respect to siCTRL treatment (data are mean ± s.d.; n = 3 technical replicates). **g,** QIBC of U2OS and RPE-1 cells immunostained for MCMBP upon siRNA treatment against DCAF12, as indicated. Lines denote medians; n ≈ 4,000 cells per condition. **h,** Quantification of QIBC in **g**, each bar indicates the median of mean intensity normalized with respect to untreated U2OS cells as 100 percent (data are mean ± s.d.; n = 2 technical replicates). **i,** MFI of cytoplasmic MCMBP under similar conditions as in **g**. **j,** Quantification of the MFI of cytoplasmic MCMBP in **i**; each bar indicates the median of mean intensity normalized with respect to untreated U2OS cells as 100 percent (data are mean ± s.d.; n = 2 technical replicates). In box plots (**c, i**), central lines denote medians, the boxes indicate the $25^{th}$ and $75^{th}$ centiles, the whiskers indicate 5 and 95 percentile values; n = 200 cells per condition. P values were calculated by ordinary one-way ANOVA with Tukey's test (**h**) or Šidák's test (**b, d, j**); **P<0.01, *P<0.1, n.s. (not significant) indicates P>0.1. (A.U. Arbitrary Units). **k,** Bone marrow whole lysates from 16-week-old DCAF12 WT and KO mice (littermates, four animals). **l,** Fetal lungs, liver, and brain whole lysates from 16-days-old DCAF12 WT and KO embryos (littermates, four animals).

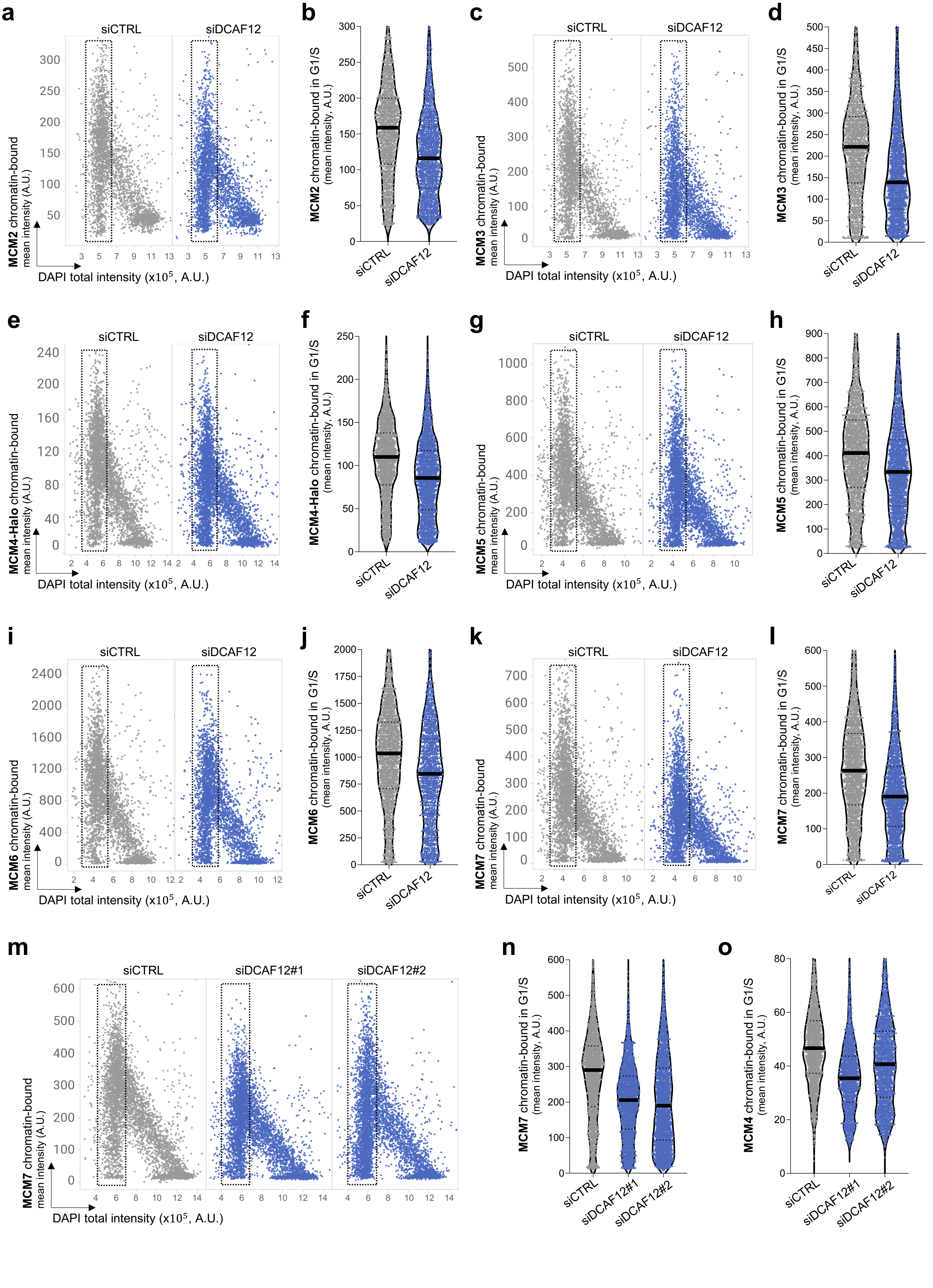


**Extended Data Fig. 4**

**Extended Data Fig. 4: Downregulation of DCAF12 results in reduced chromatin binding of all MCM subunits. a,** QIBC plots of pre-extracted U2OS cells stained for MCM2 after treatment with control siRNA or siRNA against DCAF12. DAPI counterstains nuclear DNA. n ≈ 3,500 cells per condition. Dashed boxes mark the G1/S phase stage. **b,** MFI of MCM2 in G1/S cells based QIBC in **a**. Lines denote medians; n ≈ 1,600 cells per condition. **c,** QIBC plots of pre-extracted U2OS cells stained for MCM3 after treatment as indicated. n ≈ 3,500 cells per condition. **d,** MFI of MCM3 in G1/S cells based QIBC in **c**. Lines denote medians; n ≈ 1,700 cells per condition. **e,** QIBC plots of pre-extracted U2OS cells stained for MCM4 after treatment as indicated. n ≈ 4,000 cells per condition. **f,** MFI of MCM4 in G1/S cells based QIBC in **e**. Lines denote medians; n ≈ 2,100 cells per condition. **g,** QIBC plots of pre-extracted U2OS cells stained for MCM5 after treatment as indicated. n ≈ 4,000 cells per condition. **h,** MFI of MCM5 in G1/S cells based QIBC in **g**. Lines denote medians; n ≈ 2,100 cells per condition. **i,** QIBC plots of pre-extracted U2OS cells stained for MCM6 after treatment as indicated. n ≈ 3,500 cells per condition. **j,** MFI of MCM6 in G1/S cells based QIBC in **i**. Lines denote medians; n ≈ 1,800 cells per condition. **k,** QIBC plots of pre-extracted U2OS cells stained for MCM7 after treatment as indicated. n ≈ 4,000 cells per condition. **l,** MFI of MCM7 in G1/S cells based QIBC in **k**. Lines denote medians; n ≈ 2,100 cells per condition. **m,** QIBC plots of pre-extracted U2OS cells stained for MCM7 after treatment as indicated. n ≈ 5,000 cells per condition. **n,** MFI of MCM7 in G1/S cells based QIBC in **m**. Lines denote medians; n ≈ 2,300 cells per condition. **o,** MFI of MCM4 in G1/S cells after treatment as indicated. Lines denote medians; n ≈ 1,700 cells per condition.

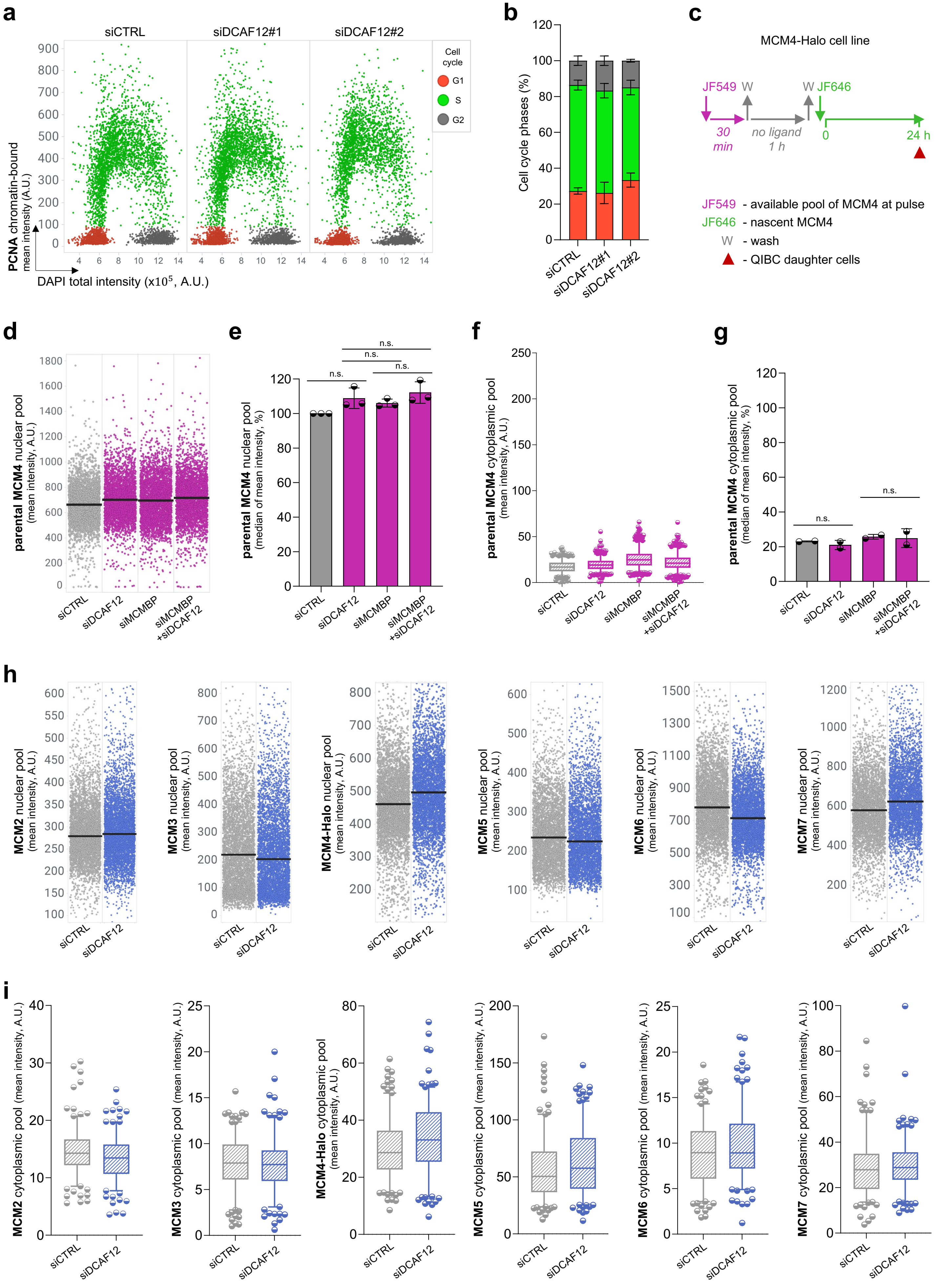


**Extended Data Fig. 5**

**Extended Data Fig. 5: Nuclear levels or localization of MCM subunits are not affected upon DCAF12 depletion. a,** QIBC plots of pre-extracted U2OS cells stained for PCNA after treatment with control siRNA or two different siRNAs against DCAF12, as indicated. DAPI counterstains nuclear DNA; n ≈ 5,000 cells per condition. **b,** Quantification of individual cell cycle phases based on QIBC in **a**; n = 2 biological replicates. **c,** Dual HaloTag labeling protocol in U2OS cells expressing endogenously tagged MCM4-Halo. **d,** QIBC plots of MCM4-Halo U2OS cells stained for parental MCM4 after indicated siRNA treatments. Lines denote medians; n ≈ 3,500 cells per condition. **e,** Quantification of QIBC plots in **d**; each bar indicates the median of mean intensity (data are mean ± s.d.; n = 3 biological replicates). **f,** MFI of cytoplasmic parental MCM4 after indicated siRNA treatments. **g,** Quantification of MFI of cytoplasmic parental MCM4; each bar indicates the median of mean intensity (data are mean ± s.d.; n = 3 biological replicates). **h,** QIBC plots of U2OS cells stained for MCM2, MCM3, MCM4-Halo, MCM5, MCM6 and MCM7, respectively; after indicated siRNA treatments. Lines denote medians; n ≈ 5,000 cells per condition (MCM2, MCM3); n ≈ 4,500 cells per condition (MCM4-Halo, MCM5, MCM6, MCM7). **i,** MFI of cytoplasmic MCM2, MCM3, MCM4, MCM5, MCM6 and MCM7, respectively; after indicated siRNA treatments. In box plots (**i**), central lines denote medians, the boxes indicate the 25$^{th}$ and 75$^{th}$ centiles, the whiskers indicate 5 and 95 percentile values; n = 200 cells per condition. P values were calculated by ordinary one-way ANOVA with Šidák's test (**e, g**); n.s. (not significant) indicates P>0.1. (A.U. Arbitrary Units).

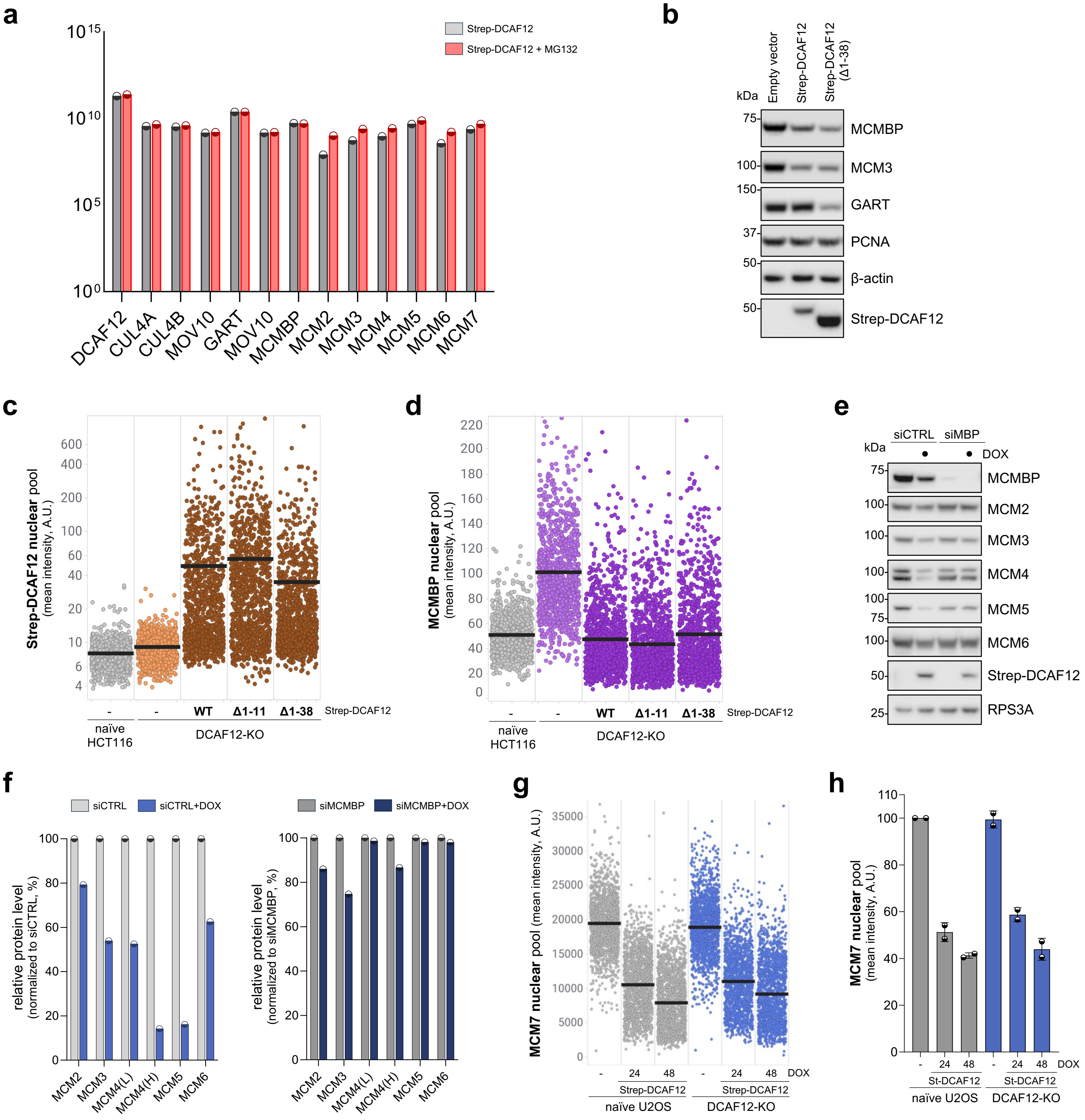


**Extended Data Fig. 6**

**Extended Data Fig. 6: Precise cellular levels of DCAF12 are critical to maintain MCMBP levels and optimal MCM equilibrium. a,** Bar graph, derived from mass spectrometry analysis of HEK293T cells co-transfected with StrepII-FLAG-tagged DCAF12 in Fig. 5f, shows the intensities of indicated proteins interacting with DCAF12 in untreated and MG132-treated samples. Cells were treated with either DMSO or 10 µM MG132 for 6 hours prior to protein purification. **b,** StrepII-FLAG-tagged DCAF12 (SF-DCAF12) or its mutant lacking NLS was inducibly expressed in HCT-116 cells using the Sleeping Beauty Transposon System. Where indicated, the expression of constructs was induced with doxycycline for 24 h. Whole-cell lysates were immunoblotted as indicated. **c, d,** QIBC plots of U2OS cells nuclei stained for StrepII-tagged DCAF12 (**c**) or MCMBP (**d**) protein after indicated DCAF12 constructs were inducibly expressed. Lines denote medians; n ≈ 1,000 cells per condition. **e**, StrepII-FLAG-tagged DCAF12 was inducibly expressed in U2OS cells using the Sleeping Beauty Transposon System. Where indicated, the expression of DCAF12 was induced with doxycycline for 24 h. Cells were transfected with control or MCMBP (MBP) targeting siRNA. Whole-cell lysates were immunoblotted as indicated. **j,** Densitometry of selected protein staining in **f; k,** QIBC plots of naïve or DCAF12 knock-out (DCAF12-KO) U2OS cells transfected with doxycycline (DOX) inducible strep-DCAF12 and immunostained for total nuclear MCM7 after 0, 24 and 48 hours of DOX treatment. Lines denote medians; n ≈ 1,500 cells per condition. **l,** Quantification of QIBC plots in **k**; each bar indicates the median of mean intensity (data are mean ± s.d.; n = 2 technical replicates).

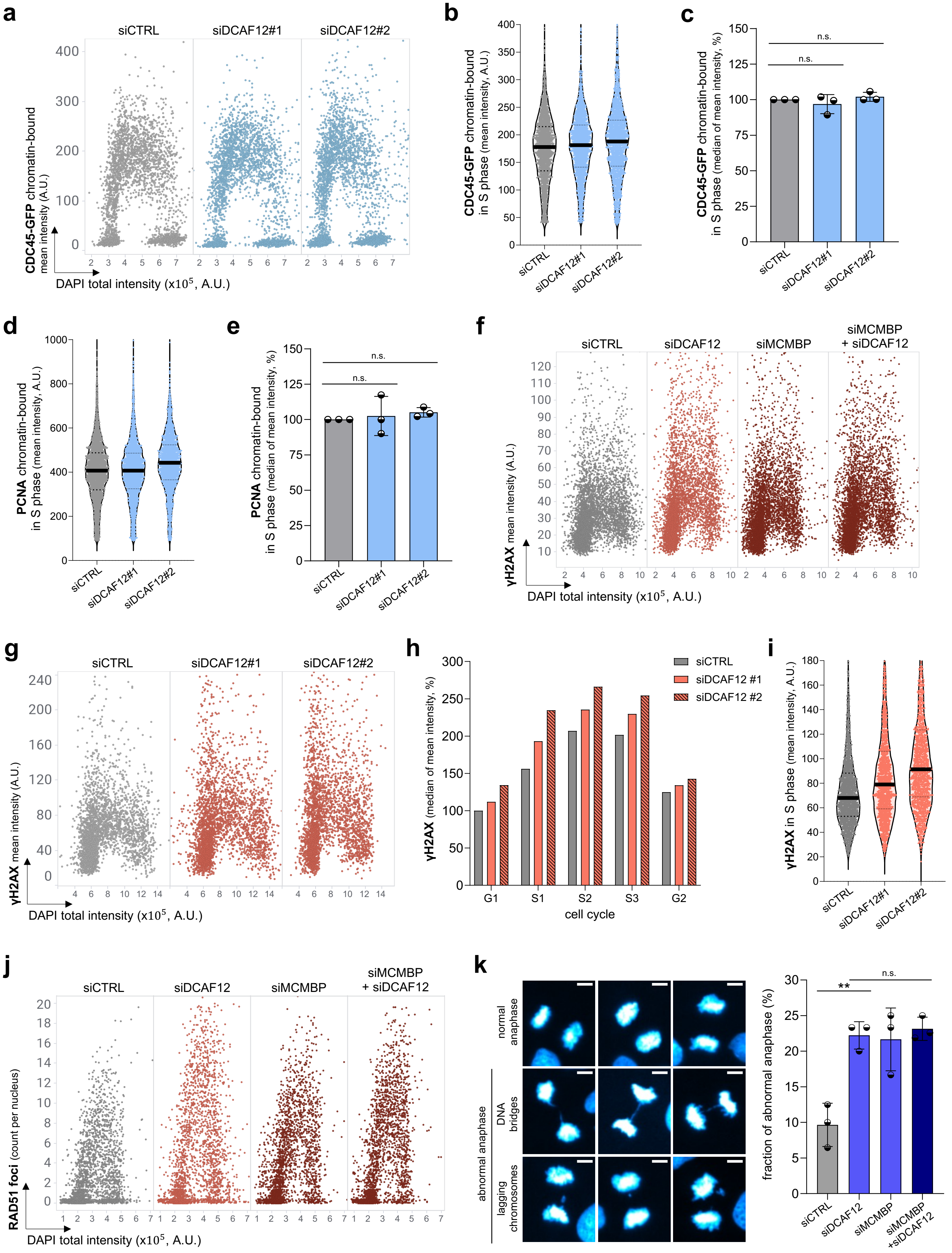


Extended Data Fig. 7

**Extended Data Fig. 7: CRL4$^{DCAF12}$-dependent regulation of MCMBP preserves genome stability. a,** QIBC plots of chromatin-bound CDC45-GFP in CDC45-GFP cells treated with control siRNA or siRNAs targeting different regions of the *DCAF12* gene (siDCAF12#1 or siDCAF12 #2), as indicated. DAPI counterstains nuclear DNA. Lines denote medians; n ≈ 3,500 cells per condition. **b,** MFI of chromatin-bound CDC45-GFP in S phase cells based on QIBC plots in **a.** Lines denote medians; n ≈ 2,000 cells per condition. **c,** Quantification of the MFI of CDC45-GFP in S phase cells based on QIBC plots in **a**. Each bar indicates the median of mean intensity normalized with respect to siCTRL as 100 percent; data are mean ± s.d.; n = 3 biological replicates. **d,** MFI of chromatin-bound PCNA in S phase cells based on QIBC in Extended Data Fig. 5a. Lines denote medians; n ≈ 2,400 cells per condition. **e,** Quantification of the MFI of PCNA in S phase cells based on QIBC in Extended Data Fig. 5a. Each bar indicates the median of mean intensity normalized with respect to siCTRL as 100 percent; data are mean ± s.d.; n = 3 biological replicates. **f,** QIBC plots of U2OS cells immunostained for γH2AX after indicated siRNA treatments. n ≈ 4,800 cells per condition. **g,** QIBC plots of U2OS cells immunostained for γH2AX after siRNA treatment as indicated. n ≈ 3,000 cells per condition. **h,** Quantification of the level of γH2AX at indicated stages of the cell cycle based on QIBC plots in **g**. Each bar indicates the median of mean intensity normalized with respect to G1 in siCTRL as 100 percent. **i,** MFI of γH2AX in S phase cells based on QIBC plots in **g**. Lines denote median, n ≈ 1,500 cells per condition. **j,** QIBC plots of RAD51 foci count per nucleus after indicated siRNA treatments, n ≈ 2,800 cells per condition. **k,** Left, representative images of anaphase anomalies such as DNA bridges and lagging chromosomes. Right, quantification of the percentage of abnormal anaphases observed after indicated siRNA treatments. Data are mean ± s.d.; n = 3 biological replicates. P values were calculated by ordinary one-way ANOVA with Šidák's test (**c, e, k** ); **P<0.01, n.s. (not significant) indicates P>0.1. (A.U. Arbitrary Units).
